# Withdraw of prophylactic antimicrobials does not change the pigs’ resistome

**DOI:** 10.1101/716597

**Authors:** Loayza-Villa Fernanda, Torres Alejandro, Zhang Lixin, Trueba Gabriel

## Abstract

The use of antimicrobials in the animal industry has increased the prevalence of antimicrobial resistant commensal bacteria in food products derived from animals, which could be associated with antimicrobial resistance in human pathogens. To reduce the influx of antibiotic resistant bacteria (and genes) to the human microbiota, restrictions on antimicrobials (in food animals) have been implemented in different countries. We investigated the impact of antimicrobial restriction in the frequency of antimicrobial resistant bacteria in pigs. No differences in antimicrobial resistance or antimicrobial resistance genes (richness or abundance) was found when we compared animals fed with and without antibiotics. Fitness costs of antimicrobial resistance in bacteria (in the field) seems to be overestimated.

## 1 Introduction

Thorough history, antimicrobials have been effective in the treatment and control of bacterial diseases and have contributed to greater life expectancy of humanity (Ferri et al., 2017). However, the emergence, spread and increasing incidence of bacteria with multiple antimicrobial resistance (AMR) has risen the concern about the use or misuse of antimicrobials (Garcia-Migura et al., 2014). Farms use around 80% of total antimicrobial production in United States (Ferri et al., 2017) and it is possible that larger proportions of antibiotics are used in animals in less industrialized countries which lack regulatory policies for antibiotic use (Ayukekbong et al., 2017).

There is a complex relationship between antimicrobial resistance in food animal microbiota and human pathogens. The use of antimicrobials in animals cause the proliferation of commensal bacteria with antimicrobial resistant genes (ARGs) which can horizontally transferred to many other bacterial species in the intestines (Forslund et al., 2013; Neu, 1992). Antimicrobial resistant commensals from farm animals can end up in food products such as meat and dairy (Van Den Bogaard and Stobberingh, 2000; Zdolec et al., 2016); these bacteria can colonize human intestines and could either become opportunistic pathogens or transfer ARGs to opportunistic pathogens (Von Wintersdorff et al., 2016; Zolec et al., 2016). This interaction between bacteria from food animals and humans has driven the creation of new policies and regulations aimed to reduce the use of antibiotics in farm animals (Chattopadhyay, 2014; Pugh, 2002)

In theory, reducing the use of antimicrobials in farms should cause a reduction in AMR bacteria commonly found in food animals and derived products (Wegener, 2003). The elimination of the selective pressure over the bacterial population should reduce the amount of AMR bacteria overtime and this process should be fast if ARGs are causing a fitness cost in commensal bacteria in the absence of antibiotics (Andersson and Hughes, 2010, 2012). However, many experiments in which animals were deprived of antibiotics as growth promotors, showed high levels of antibiotic resistance in numerically dominant *Escherichia coli* (Ahmed et al., 2017; Mathew et al., 1998) or high relative abundance of resistance genes (Pakpour et al., 2012). More importantly, resistant genes, multi-resistant and numerically dominant bacteria have been found associate to animal production in organic farms too (Gerzova et al., 2015; Österberg et al., 2016; Mollenkopf et al., 2018). In the present study we investigated the effect of the removal of antibiotics administered as prophylactics (higher antibiotic doses than for growth promotion). We analyzed phenotypic resistance in coliforms and microbiota resistome.

## 2 Material and methods

### 2.1 Animals

A random, balanced double-blind study was conducted in two generations of pigs. Twenty healthy female 70d piglets were selected and separated in two pens with 10 piglets each. One group was fed with antimicrobial additives (group A) and, 10 without antimicrobial additives in feed (group B). Treatments were maintained during growth, sexual maturity and pregnancy. Once sows farrowed, piglets were weaned and placed in two separate pens for group A (n=32) and B (n=32) respectively and continued with treatments of their respective mothers. All the experimental procedures were approved by the Ethics Committee for Animal Research of San Francisco de Quito University. Vaccines were administered to all animals and the antimicrobial treatment was administered under veterinarian supervision to animals that have any diagnosed infection. Antimicrobial additives used are described in Table 1.

**Table 1.** Antimicrobial additives used in pigs farm as prophylactics in group A.

| Growth phase | Age (days) | Antimicrobial | Dose, ppm | Administration via |
| --- | --- | --- | --- | --- |
| 0 | 21 - 28 | Tilmicosin | 200 | Feed |
|  |  | Colistin | 40 | Feed |
| 1 | 29 - 34 | Tiamulin | 150 | Feed |
|  |  | Chlortetracycline | 450 | Feed |
| 2 | 35 - 45 | Tiamulin | 150 | Feed |
|  |  | Chlortetracycline | 450 | Feed |
| 3 | 45 - 70 | Tiamulin | 150 | Feed |
|  |  | Chlortetracycline | 450 | Feed |
| 4 | 70 - 85 | Chlortetracycline | 450 | Feed |
| 5 | 123 - 139 | Chlortetracycline | 450 | Feed |
| 2 | 37 - 40 | Trimetoprim-sulfamethoxazole | 25mg/Kg/PV | Water |
| 3 | 45 - 47 | Doxicycline | 10mg/Kg/PV | Water |

### 2.2 Samples and phenotypic analysis

Rectal swabs were taken from sows and 5 randomly selected piglets from each litter during one productive cycle (Figure 1). During weaning and fattening phases, each pen grouped 32 piglets. Pig density was 0,45 m^2^/pig in weaning phase and 0,90-1,0m^2^/pig at fattening phase. Animals from each group were monitored for 170 days (Figure 1). The type and antimicrobial concentrations in feed changed overtime and have been used routinely in the farm for the two previous years (table 1).

**Figure 1.**
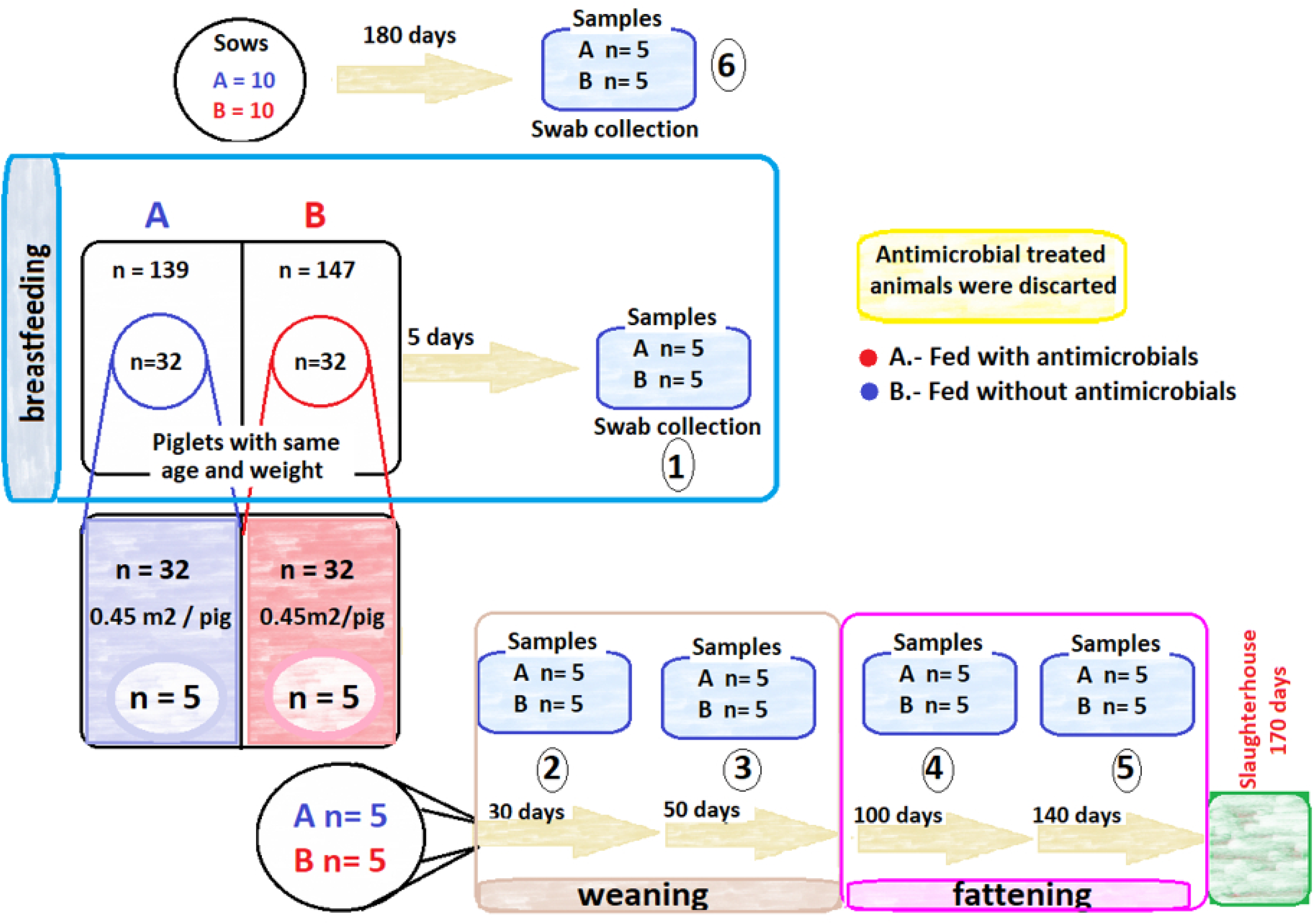
Work flow for a productive pig’s cycle of each treatment group A. fed with antimicrobials. B. fed without antimicrobials. 10 female pigs (sows at 70 days) were randomly selected for each treatment. All farrowed piglets were maintained under the same treatment group until day 2l(Breastfeeding phase). A homogeneous group of 32 piglets (similar age and weight) within each treatment group were selected to form a weaning phase pen (since 22 days to 70 days) that was conserve until the slaughter phase (170 days). Rectal svvabs samples were collected at 1.- 5 days; 2.- 30days; 3.- 50days; 4.- 100 days; 5.- 140 days. A sample from sows were taken at l 80days (6.)

Swabs were maintained on ice for transportation to the lab facilities within 2h after collection. For molecular analysis, samples were frozen at -80°C. Intestinal coliforms were used as microbial indicator of phenotypic resistance. Swabs were eluted in 1mL of sterile phosphate buffered saline solution (PO), 0.1mL of this solution with be serial diluted in 0.9mL of PO until 10-3. Then, 0.1mL of dilution of the sample was plated onto the surface of MacConkey Agar (MKL) with and without antimicrobials (Table 2).

**Table 2.**
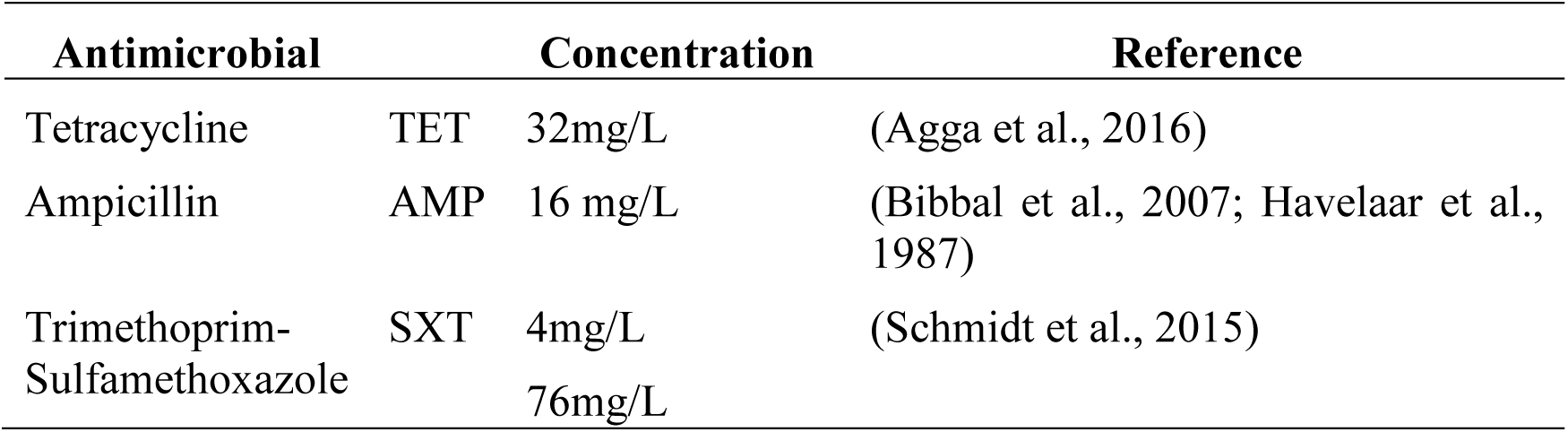
Antimicrobials used as supplements to MacConkey Lactose (MKL) culture media.

We estimated the ratio of resistant coliforms by counting the number of colonies in plates with antimicrobials divided by the colonies in MKL plate without antimicrobials.

### 2.3 Antimicrobial susceptibility test

One lactose fermenting (coliform) colony from each plate was isolated and stored at -80C in TSB with 10% of glycerol. Antimicrobial susceptibility tests were performed with Bauer Kirby test following CLSI guidelines, on random selected strains using AMP ampicillin (10μg), TET tetracycline (30μg), SXT trimethoprim-sulfamethoxazole (1.25/23.75μg), GEN gentamycin (10μg), AMC amoxicillin-clavulanic ac. (20/10μg), CIP ciprofloxacin (5μg), CHLOR chloramphenicol (30μg) and COX ceftriaxone (30μg) as representatives of the most used families of antibacterial drugs used in health care (Eisenberg et al., 2012; Kozak et al., 2009).

### 2.4 Molecular analysis

DNA from swabs taken for each pig were isolated using MO BIO Power Soil DNA Isolation Kit (MO BIO, 12888-100) using swab dilution material in bead solution buffer and following manufacturer instructions. Quality and quantity were evaluated using nanodrop (Thermo Scientific) and Qubit dsDNA HS (Thermo Fisher Scientific, Oregon, USA)

From sows’ samples, *mcr*-1 PCR amplification were performed used the conditions described previously (Liu et al., 2016). One pooled sample from each sampling point (6 from A and 6 from B group) were analyzed in duplicate with high throughput qPCR. WaferGen SmartChip Real-time PCR system was performed to detect 384 genes, 338 are informative for AR genes or MGE. Primers for these genes and associated HT-qPCR assay were designed, used, and validated in the previous studies (Guo et al., 2018; Looft et al., 2012; Su et al., 2015; Zhu et al., 2017), and primer set was update recently (Stedtfeld et al., 2018). The genetic richness was defined as the number of AMR genes found in a niche.

### 2.5 Statistical analysis

All collected data were registered in MS EXCEL software and descriptive and inferential statistics analysis were performed in INFOSTAT (Statistic Software Vs 2017). The impact of antimicrobial restriction on coliforms count, on the susceptibility patterns of isolates and animal performance were compared with T test and Chi Square respectively (p≤0.05). HT-qPCR data was analyzed according to previously established methods (Looft et al., 2012; Muurinen et al., 2017). Specifically, ΔΔCT method was used to normalize and calculate the fold change. Moreover, the relative abundance of ARG was calculated with normalization to the universal 16S rRNA; estimated from the Ct value with a conservative threshold Ct of 30 as the gene copy detection limit due to the lack of quantification curves for each 384 primer sets used. Calculated data represent the copy number per 16S rRNA gene copy. QIUcore Omics Explore 3.4 software were used to show heat maps.

## 3 Results

Based on coliform counts in MKL with and without antimicrobials, AMR ratios of resistant coliforms to overall coliforms were calculate and are shown in Table 3. The resistance ratios for tetracycline were higher than that for trimethoprim-sulfamethoxazole or ampicillin. No significant differences between treatment groups for any antibiotic was found (p: 0.434, 0.722, 0,763 respectively)

**Table 3.** Total count of coliform colony forming units (CFU) in Mac Conkey lactose without antimicrobials. The average and standard deviation (SD) is shown for each treatment. Antimicrobial resistance ratios for ampicillin (AMP), cotrimoxazole (SXT) and Tetracycline (TET) were calculate using the total count of coliform colony forming units in Mac Conkey Lactosa plates with antimicrobials divided by the total count of coliform in Mac Conkey Lactosa without antimicrobials.

| AGE (days) | Treatment A |  |  |  |  | Treatment B |  |  |  |  |
| --- | --- | --- | --- | --- | --- | --- | --- | --- | --- | --- |
|  | (AVERAGE) | SD | AMP | SXT | TET | 2,34E+07 | SD | AM | SXT | TET |
| 5 | 1,73E+07 | 1,15E+07 | 0,48 | 1,22 | 2,15 | 5,15E+04 | 1,10E+13 | 0,39 | 1,55 | 2,38 |
| 30 | 6,71E+05 | 1,01E+06 | 0,78 | 0,56 | 0,66 | 8,90E+05 | 7,92E+14 | 0,98 | 0,43 | 0,97 |
| 50 | 9,17E+05 | 8,70E+05 | 0,26 | 0,48 | 0,83 | 7,40E+04 | 8,41E+14 | 0,48 | 0,48 | 1,01 |
| 100 | 2,46E+05 | 1,68E+05 | 0,38 | 0,94 | 0,99 | 2,65E+05 | 1,16E+13 | 0,48 | 1,48 | 0,72 |
| 140 | 1,08E+05 | 1,68E+05 | 0,34 | 0,32 | 1,05 |  | 2,55E+14 | 0,25 | 0,14 | 1,2 |

Antimicrobial susceptibility tests for 537 randomly selected strains (A=266 and B= 271) showed general resistance to ampicillin (n= 397; 73.9%), amoxicillin-clavulanic ac. (n=188; 35%), tetracycline (n=434; 81.1%), trimethoprim-sulfamethoxazole (n=301; 56.1%), gentamycin (n=125; 23.3%), ciprofloxacin (n=71; 13.2%), chloramphenicol (n= 174; 32.4%) and ceftriaxone (n=77; 14.3%) were detected. There were no significative differences (p ≥ 0.05) between treatment groups neither in sows nor in piglets (in nursing or fattening phases) (Table 4). Strains with resistance to 3 or more antimicrobials were considered as multidrug resistant phenotype (MDR) (n= 354; 65.9%).

**Table 4.**
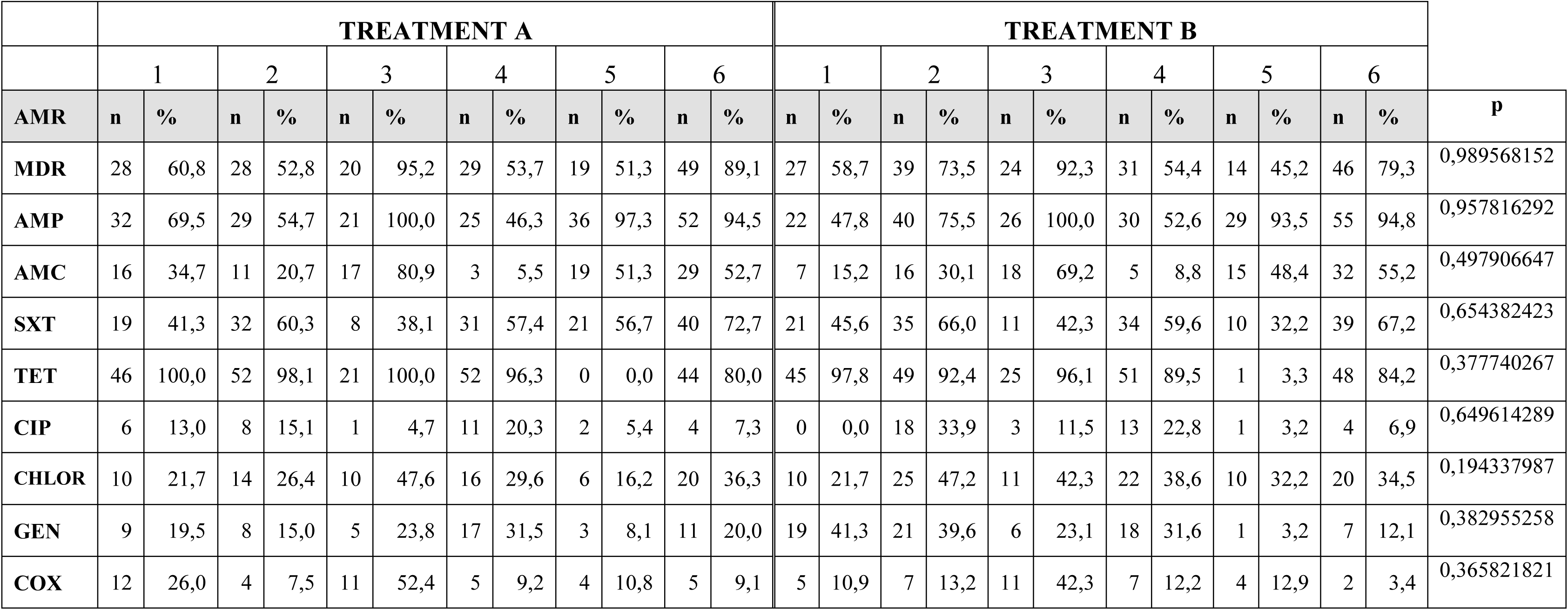
Antimicrobial susceptibility test from coliform isolated. Strains are classified by sampling period (1.-5days; 2-30days; 3.-50 days; 4.-100 days; 5.-140 days) and treatment group (A.-with antimicrobials; B.-without antimicrobials) (p = 0,77). p was calculated based on sampling period comparison. Strain with more than 2 resistances was count as multidrug resistant (MDR). AMP ampicillin (10μg), TET tetracycline (30μg), SXT trimethoprim-sulfamethoxazole (1.25/23.75μg), GEN gentamycin (10μg), AMC amoxicillin-clavulanic ac. (20/10μg), CIP ciprofloxacin (5μg), CHLOR chloramphenicol (30μg) and COX ceftriaxone (30μg) were used to perform the antimicrobial susceptibility test.

The antimicrobial resistance richness was not different (p≥ 0.05) in animals within group neither between groups (Figure 3) and the abundance of resistant genes decreased overtime, although tetracycline resistance genes and mobile genetic elements (MGE) remained stable. Among the most abundant genes detected were aminoglycoside resistance and MGE (Tp614, IS613, tnpA, int1-a-marko, intl2, intI1F165_ clinical, pBS228-IncP-1, trb-C, IS26, IS256, IS6100, IS91). These MGEs could be responsible for the transference of resistance genes among microbiota species; tet(32) was detected in all samples. Colistin resistant gene were found too but the frequency was low and was reported within “Other” category. Furthermore, a PCR amplification were performed on sows’ samples at the beginning of this study. *mcr*-1 was amplified in 19 from 20 of these samples.

**Figure 2.**
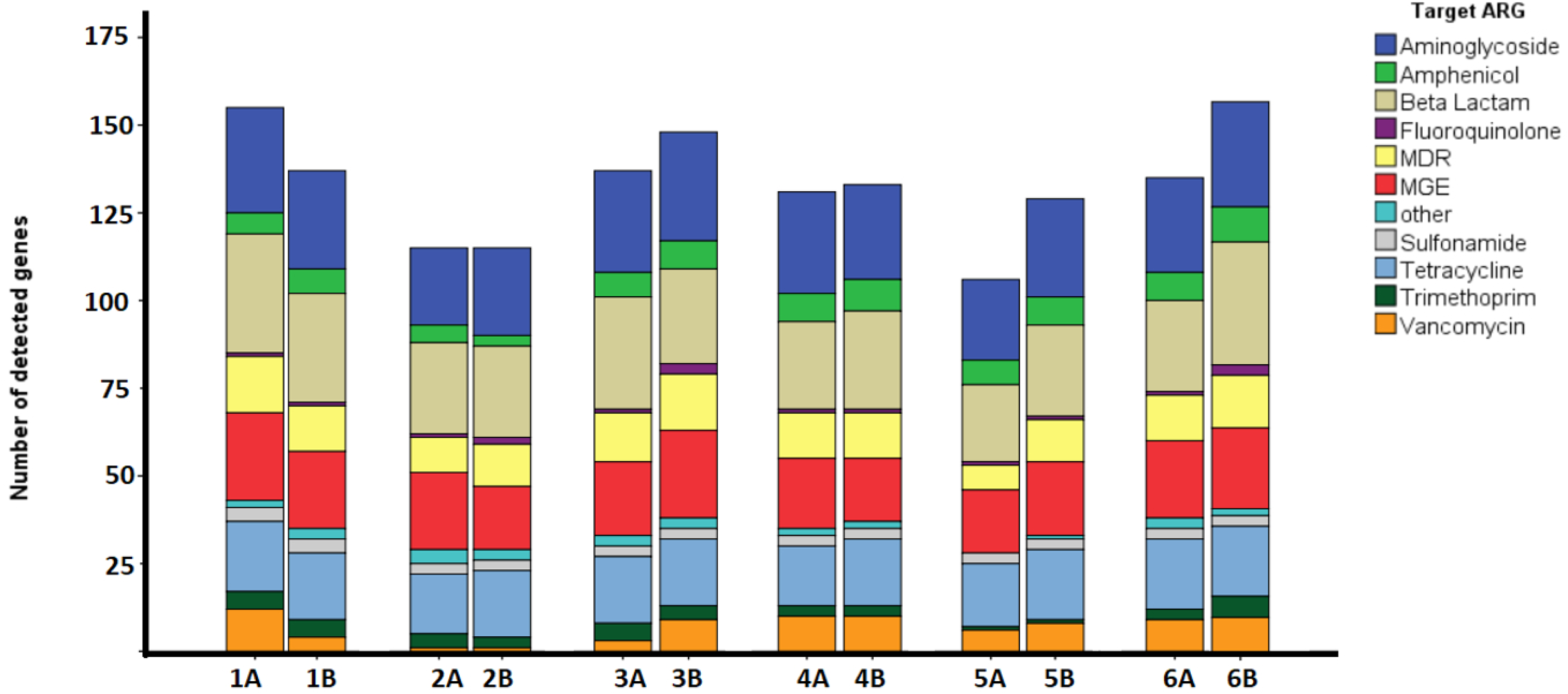
Antimicrobial resistance genes (ARG) richness. Number of genes of each class of antimicrobial target. In columns there are assigned the growing phase ofpiglets (1.- 5 days, 2.- 30 days; 3.- 50 days, 4.- 100 days; 5.- 140 days) and sows (6.- 180days). Animals feeding antimicrobials were identified as A and animal without antimicrobial additives were identified with B.

**Figure 3.**
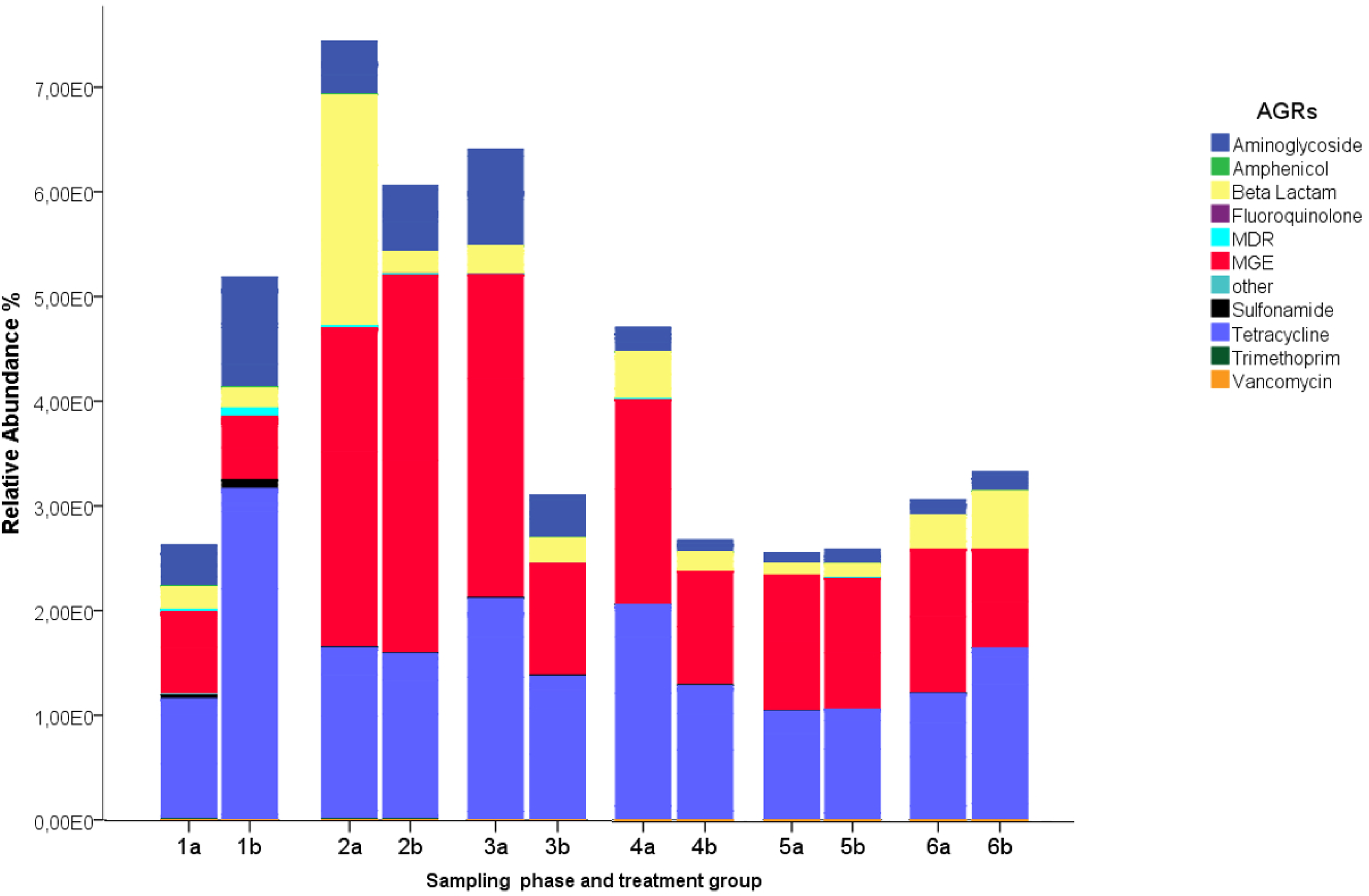
Relative abundance (RA%) of antimicrobial resistance genes grouped by samplingphase and treatment group (A.-Pigs fed antibiotics; B.-Pigs not feed antibiotics); 1.- 5 days; 2.- 30 days; 3.- 50 days; 4.- 100 days; 5.-140 days; 6 Sows 180 days)

**Figure 4.**
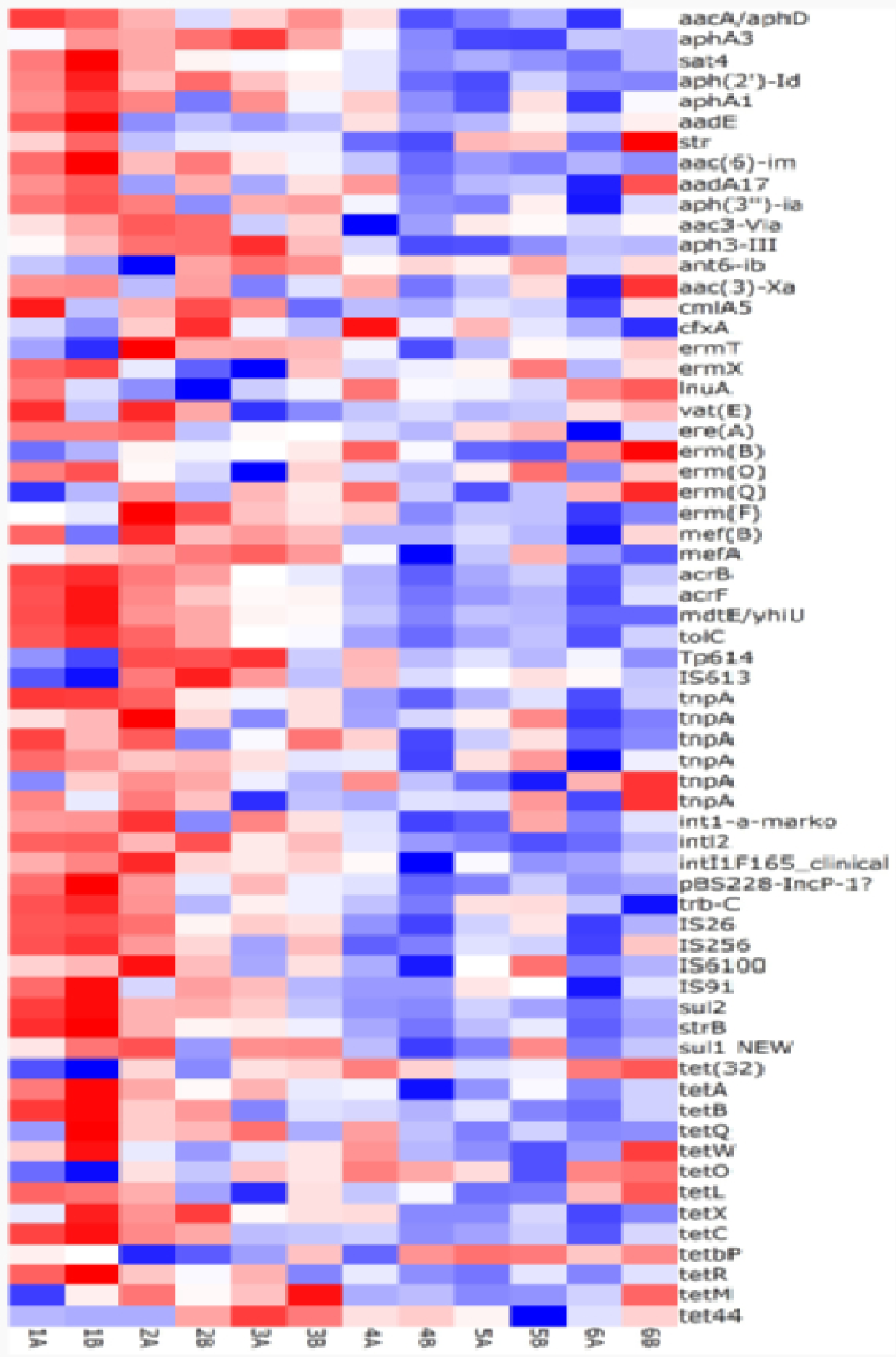
Relative abundance (RA%) of ARG genes grouped by sampling phase and heatment group (A.-Pigs feeding with antibiotics; B.-Pigs feeding without antibiotics); 1. -5d; 2. -30d; 3.- 50d; 4.- 100d; 5.- 140d; 6 Sows at 180d). Genes with CT <30 were excluded, the heat map represents only genes detected in at least one sample. hl red are shown genes with higher relative abundance, in blue the less abundant genes.

A Sperman correlation test was performed on QIUcore Omics Explore 3.4 software (Supl. 16) showed no difference of ARG relative abundance profiles between samples collected during growth phases. Pigs at day 30, showed a higher ARG relative abundance (although not statistically significant) than pigs at day 5, however there was no statistical difference between groups (figure 3). However, this high ARG relative abundance declined overtime.

## 4 Discussion

In this study, we found that antimicrobial restriction (during 2 generations of pigs) had no significant impact in antibiotic resistance of intestinal coliforms. We hypothesized that the absence of antimicrobials in the diet (during two generations of animals) will cause antimicrobial sensible bacteria to outgrow resistant ones. However, we did not find significant differences (α ≥ 0.05) in the total number of resistant coliforms nor did we find differences in resistance gene abundance or diversity associated with antibiotic additives. Similar results, in pathogens and commensals, have been reported previously (Ahmed et al. 2017; Miranda et al., 2009; Cho et al., 2007; Sato et al., 2004). The lack of differences in ARG relative abundance between groups of pigs feeding with and without antimicrobials was also reported (Gerzova et al., 2015). Other studies found that *E. coli* isolated from sows with different levels of antimicrobials in feed, had some differences in antibiotic resistance genes but similar antibiotic susceptibility phenotypes (Mazurek et al., 2014; Looft et al., 2014); other studies, however, showed an important decrease in resistance bacterial isolates or reduced resistant gene abundance after antibiotic removal (Österberg et al., 2016; Looft et al., 2012; Mathew et al, 1998). These discrepancies may be due to differences in fitness cost of the different plasmids over different bacterial populations. Some authors suggest that longer periods of antibiotic restriction are required in order to observe some changes in the relative abundance of antimicrobial resistant genes (Pakpour et al., 2012), however, long time (> 10 years) reductions in antibiotic use have resulted in almost null reduction of antibiotic resistance in Enterobacteriacea from food-animals (Danish Integrated Antimicrobial Resistance Monitoring, Research Programme, 2017). These findings are also in agreement with the notion that the resistome tends to persist over time even in the absence of selective pressure (Lehtinen et al., 2017).

The resistance genes with higher relative abundance were against tetracyclines, β lactams, and aminoglycosides (Figure 3). These phenotypes have been commonly reported in pig farms (Österberg et al., 2016). Genes associated with tetracycline resistance have been detected in pigs feeding with or without medicated feed which concords with the notion tetracycline resistance genes are common in swine intestinal resistome (Koga et al., 2015; Looft et al., 2012; Kazimierczak et al., 2009; Mathew et al, 1998). β-lactamase or aminoglycoside resistance genes were detected despite these antimicrobials were absent in the diet. Similar observations have been reported by Looft et al. (2012) and were confirmed by the same research group in 2014. The mobile genetic elements markers, that showed an important relative abundance, may explain the abundance of these genes (Looft et al., 2014).

Our results may indicate that ARGs are not causing any fitness reduction in the bacterial population in the intestine. Under laboratory conditions it has been shown that fitness costs, associated a plasmid, could be transformed in fitness advantages after 420 generations (Dionisio et al., 2005); and the advantageous plasmid could improve the fitness of bacterial hosts never exposed to this plasmid (Dionisio et al., 2005).

MGEs are important actors in antimicrobial resistance spread (Jansen and Aktipis, 2014) and in this study, we observed a higher relative abundance of MGEs in piglets at 30 day of age which coincides with the weaning period. This phenomenon may be linked to gut microbiome stress due to changes in diet (Frese et al., 2015; Gresse et al., 2017; Isaacson and Kim, 2012); bacteria under stress may turn on their S.O.S response which increases mobilization of MGEs (Shapiro, 2015; Beaber et al., 2014). This perturbation of intestinal microbiota has been associated with higher activity of MGEs and abundance of resistance genes in piglets (Twiss et al., 2005)

Our study had some limitations such as housing the two groups of animals in the same barn. The environment may be saturated with resistant clones and our results may be driven by this environmental exposure. However, this limitation doesn’t invalidate our findings as we wanted to investigate whether antibiotic-sensitive bacteria in the intestine are a better fit to grow in the intestines in the absence of antibiotics (Aarestrup, 2015; Wasyl et al., 2012). Finally, the withdraw of antibiotics in this setting did not have any repercussion in the growth or the health of these animals.

## 5 Conclusion

Our observations suggest that antibiotic restriction is not enough to reduce the numbers of antibiotic-resistant bacteria in the gastrointestinal tract of food animals and their products. The maintenance of antibiotic resistance in the absence of antibiotic pressure is not easily explained, there are many evolutionary factors that are not fully understood and require additional research.

## 6 Authors contribution

GT Funding acquisition, Farm access agreement and project conceptualization

FL Experimental design, field and lab techniques, sampling procedures, data register and analysis, write and edition of publication.

AT experimental design, farm permissions and animal welfare.

LZ Molecular methodology, technical support and data analysis.

LZ and GT manuscript edition

## 9 Acknowledgments

We thank all technicians from laboratory of veterinarian diagnostics in ecuadorian food animal industry, specially Marcela Coba and Juan Castellanos. We appreciate the helpful support of Edgar Velarde and Diego Velasquez and all people in Microbiology Institute at San Francisco de Quito University.

## Reference

Aarestrup, F. M. (2015). The livestock reservoir for antimicrobial resistance: a personal view on changing patterns of risks, effects of interventions and the way forward. Philos. Trans. R. Soc. Lond. B. Biol. Sci. 370, 20140085. doi:10.1098/rstb.2014.0085

Ahmed, S., Olsen, J. E., and Herrero-Fresno, A. (2017). The genetic diversity of commensal *Escherichia coli* strains isolated from non- antimicrobial treated pigs varies according to age group. PLoS One 12, 1–18. doi:10.1371/journal.pone.0178623.

Andersson, D. I., and Hughes, D. (2010). Antibiotic resistance and its cost: is it possible to reverse resistance? Nat. Rev. Microbiol. 8, 260–271. doi:10.1038/nrmicro2319.

Andersson, D. I., and Hughes, D. (2012). Evolution of antibiotic resistance at non-lethal drug concentrations. Drug Resist. Updat. 15, 162–72. doi:10.1016/j.drup.2012.03.005.

Ayukekbong, J. A., Ntemgwa, M., and Atabe, A. N. (2017). The threat of antimicrobial resistance in developing countries: causes and control strategies. Antimicrob. Resist. Infect. Control 6, 47. doi:10.1186/s13756-017-0208-x.

Beaber, J. W., Hochhut, B., and Waldor, M. K. (2004). SOS response promotes horizontal dissemination of antibiotic resistance genes. Nature 427, 72–74. doi:10.1038/nature02241

Bibbal, D., Dupouy, V., Ferré, J. P., Toutain, P. L., Fayet, O., Prè, M. F., et al. (2007). Impact of three ampicillin dosage regimens on selection of ampicillin resistance in Enterobacteriaceae and excretion of *bla*TEM genes in swine feces. Appl. Environ. Microbiol. 73, 4785–4790. doi:10.1128/AEM.00252-07.

Chattopadhyay, M. K. (2014). Use of antibiotics as feed additives: a burning question. Front. Microbiol. 5, 334. doi:10.3389/fmicb.2014.00334.

Cho, S., Fossler, C. P., Diez-Gonzalez, F., Wells, S. J., Hedberg, C. W., Kaneene, J. B., et al. (2007). Antimicrobial Susceptibility of Shiga Toxin-Producing Escherichia coli Isolated from Organic Dairy Farms, Conventional Dairy Farms, and County Fairs in Minnesota. Foodborne Pathog. Dis. 4, 178–186. doi:10.1089/fpd.2006.0074

Dionisio, F., Conceição, I. C., Marques, A. C. R., Fernandes, L., and Gordo, I. (2005). The evolution of a conjugative plasmid and its ability to increase bacterial fitness. Biol. Lett. 1, 250–252. doi:10.1098/rsbl.2004.0275.

Eisenberg, J. N. S., Goldstick, J., Cevallos, W., Trueba, G., Levy, K., Scott, J., et al. (2012). In-roads to the spread of antibiotic resistance: regional patterns of microbial transmission in northern coastal Ecuador. J. R. Soc. Interface 9, 1029–1039. doi:10.1098/rsif.2011.0499.

Ferri, M., Ranucci, E., Romagnoli, P., and Giaccone, V. (2017). Antimicrobial resistance: A global emerging threat to public health systems. Crit. Rev. Food Sci. Nutr. 57, 2857–2876. doi:10.1080/10408398.2015.1077192.

Forslund, K., Sunagawa, S., Kultima, J. R., Mende, D. R., Arumugam, M., Typas, A., et al. (2013). Country-specific antibiotic use practices impact the human gut resistome. Genome Res. 23, 1163–1169. doi:10.1101/gr.155465.113.23

Frese, S. A., Parker, K., Calvert, C. C., and Mills, D. A. (2015). Diet shapes the gut microbiome of pigs during nursing and weaning. Microbiome 3, 1–10. doi:10.1186/s40168-015-0091-8.

Garcia-Migura, L., Hendriksen, R. S., Fraile, L., and Aarestrup, F. M. (2014). Antimicrobial resistance of zoonotic and commensal bacteria in Europe: The missing link between consumption and resistance in veterinary medicine. Vet. Microbiol. 170, 1–9. doi:10.1016/j.vetmic.2014.01.013.

Gerzova, L., Babak, V., Sedlar, K., Faldynova, M., Videnska, P., Cejkova, D., et al. (2015). Characterization of antibiotic resistance gene abundance and microbiota composition in feces of organic and conventional pigs from four EU countries. PLoS One 10, 1–10. doi:10.1371/journal.pone.0132892.

Gresse, R., Chaucheyras-Durand, F., Fleury, M. A., Van de Wiele, T., Forano, E., and Blanquet-Diot, S. (2017). Gut Microbiota dysbiosis in postweaning piglets: understanding the keys to health. Trends Microbiol. 25, 851–873. doi:10.1016/j.tim.2017.05.004.

Guo, X., Stedtfeld, R. D., Hedman, H., Eisenberg, J. N. S., Trueba, G., Yin, D., et al. (2018). Antibiotic Resistome Associated with Small-Scale Poultry Production in Rural Ecuador. Environ. Sci. Technol. 52, 8165–8172. doi:10.1021/acs.est.8b01667.

Havelaar, A. H., During, M., and Versteegh, J. F. M. (1987). Ampicillin-dextrin agar medium for the enumeration of *Aeromonas* species in water by membrane filtration. J. Appl. Bacteriol. 62, 279–287. doi:10.1111/j.1365-2672.1987.tb02410.x.

Isaacson, R., and Kim, H. B. (2012). The intestinal microbiome of the pig. Anim. Heal. Res. Rev. 13, 100–109. doi:10.1017/S1466252312000084

Jansen, G., and Aktipis, C. A. (2014). Resistance Is Mobile: The Accelerating Evolution of Mobile Genetic Elements Encoding Resistance. J. Evol. Med. 2, 1–3. doi:10.4303/jem/235873.

Kazimierczak, K. A., Scott, K. P., Kelly, D., and Aminov, R. I. (2009). Tetracycline Resistome of the Organic Pig Gut. Appl. Environ. Microbiol. 75, 1717–1722. doi:10.1128/AEM.02206-08.

Koga, V. L., Scandorieiro, S., Vespero, E. C., Oba, A., De Brito, B. G., De Brito, K. C. T., et al. (2015). Comparison of Antibiotic Resistance and Virulence Factors among *Escherichia coli* Isolated from Conventional and Free-Range Poultry. Biomed Res. Int. 2015. doi:10.1155/2015/618752.

Kozak, G. K., Boerlin, P., Janecko, N., Reid-Smith, R. J., and Jardine, C. (2009). Antimicrobial resistance in *Escherichia coli* isolates from swine and wild small mammals in the proximity of swine farms and in natural environments in Ontario, Canada. Appl. Environ. Microbiol. 75, 559–566. doi:10.1128/AEM.01821-08.

Lehtinen, S., Blanquart, F., Croucher, N. J., Turner, P., Lipsitch, M., and Fraser, C. (2017). Evolution of antibiotic resistance is linked to any genetic mechanism affecting bacterial duration of carriage. Proc. Natl. Acad. Sci. U. S. A. 114, 1075–1080. doi:10.1073/pnas.1617849114

Liu, Y. Y., Wang, Y., Walsh, T. R., Yi, L. X., Zhang, R., Spencer, J., et al. (2016). Emergence of plasmid-mediated colistin resistance mechanism MCR-1 in animals and human beings in China: A microbiological and molecular biological study. Lancet Infect. Dis. 16, 161–168. doi:10.1016/S1473-3099(15)00424-7

Looft, T., Johnson, T. A., Allen, H. K., Bayles, D. O., Alt, D. P., Stedtfeld, R. D., et al. (2012). Infeed antibiotic effects on the swine intestinal microbiome. Proc. Natl. Acad. Sci. 109, 1691–1696. doi:10.1073/pnas.1120238109.

Looft, T., Allen, H. K., Cantarel, B. L., Levine, U. Y., Bayles, D. O., Alt, D. P., et al. (2014). Bacteria, phages and pigs?: the effects of in-feed antibiotics on the microbiome at different gut locations. ISME J. 8, 1566–1576. doi:10.1038/ismej.2014.12

Mathew, A. G., Upchurch, W. G., and Chattin, S. E. (1998). Incidence of Antibiotic Resistance in Fecal *Escherichia coli* Isolated from Commercial Swine Farms. J. Anim. Sci. 76, 429–434.

Mazurek, J., Bok, E., Pusz, P., Stosik, M., and Baldy, K. (2014). Phenotypic and genotypic characteristics of antibiotic resistance of commensal Escherichia coli isolates from healthy pigs. 211–218. doi:10.2478/bvip-2014-0031.

Miranda, J. M., Mondragón, A., Vázquez, B. I., Fente, C. A., Cepeda, A., and Franco, C. M. (2009). Influence of farming methods on microbiological contamination and prevalence of resistance to antimicrobial drugs in isolates from beef. Meat Sci. 82, 284–288. doi:10.1016/j.meatsci.2009.01.020.

Mollenkopf, D. F., De Wolf, B., Feicht, S. M., Cenera, J. K., King, C. A., van Balen, J. C., et al. (2018). *Salmonella* spp. and extended-spectrum cephalosporin-resistant *Escherichia coli* frequently contaminate broiler chicken transport cages of an organic production company. Foodborne Pathog. Dis. 15, 583–588. doi:10.1089/fpd.2017.2390.

Muurinen, J., Karkman, A., and Virta, M. (2017). “High Throughput Method for Analyzing Antibiotic Resistance Genes in Wastewater Treatment Plants,” in Antimicrobial Resistance in Wastewater Treatment Processes (Hoboken, NJ, USA: John Wiley & Sons, Inc.), 253–262. doi:10.1002/9781119192428.ch14.

Neu, H. C. (1992). The crisis in antibiotic resistance. Science 257, 1064–1073. doi:10.1126/science.257.5073.1064.

Österberg, J., Wingstrand, A., Jensen, A. N., Kerouanton, A., Cibin, V., Barco, L., et al. (2016). Antibiotic resistance in Escherichia coli from pigs in organic and conventional farming in four european countries. PLoS One 11, 1–12. doi:10.1371/journal.pone.0157049

Pakpour, S., Jabaji, S., and Ch??nier, M. R. (2012). Frequency of Antibiotic Resistance in a Swine Facility 2.5 Years After a Ban on Antibiotics. Microb. Ecol. 63, 41–50. doi:10.1007/s00248-011-9954-0.

Pugh, D. M. (2002). The EU precautionary bans of animal feed additive antibiotics. Toxicol. Lett. 128, 35–44.

Sato, K., Bartlett, P. C., Kaneene, J. B., and Downes, F. P. (2004). Comparison of prevalence and antimicrobial susceptibilities of *Campylobacter spp*. isolates from organic and conventional dairy herds in Wisconsin. Appl. Environ. Microbiol. 70, 1442–1447. doi:10.1128/AEM.70.3.1442-1447.2004.

Shapiro, R. S. (2015). Antimicrobial-Induced DNA Damage and Genomic Instability in Microbial Pathogens. PLOS Pathog. 11, e1004678. doi:10.1371/journal.ppat.1004678

Stedtfeld, R. D., Guo, X., Stedtfeld, T. M., Sheng, H., Williams, M. R., Hauschild, K., et al. (2018). Primer set 2.0 for highly parallel qPCR array targeting antibiotic resistance genes and mobile genetic elements. FEMS Microbiol. Ecol. 94. doi:10.1093/femsec/fiy130.

Su, J.-Q., Wei, B., Ou-Yang, W.-Y., Huang, F.-Y., Zhao, Y., Xu, H.-J., et al. (2015). Antibiotic Resistome and Its Association with Bacterial Communities during Sewage Sludge Composting. Environ. Sci. Technol. 49, 7356–7363. doi:10.1021/acs.est.5b01012.

Twiss, E., Coros, A. M., Tavakoli, N. P., and Derbyshire, K. M. (2005). Transposition is modulated by a diverse set of host factors in Escherichia coli and is stimulated by nutritional stress. Mol. Microbiol. 57, 1593–1607. doi:10.1111/j.1365-2958.2005.04794.x.

Van Den Bogaard, A. E., and Stobberingh, E. E. (2000). Epidemiology of resistance to antibiotics: Links between animals and humans. Int. J. Antimicrob. Agents 14, 327–335. doi:10.1016/S0924-8579(00)00145-X.

Von Wintersdorff, C. J. H., Penders, J., Van Niekerk, J. M., Mills, N. D., Majumder, S., Van Alphen, L. B., et al. (2016). Dissemination of antimicrobial resistance in microbial ecosystems through horizontal gene transfer. Front. Microbiol. 7, 1–10. doi:10.3389/fmicb.2016.00173.

Wasyl, D., Hasman, H., Cavaco, L. M., and Aarestrup, F. M. (2012). Prevalence and characterization of cephalosporin resistance in nonpathogenic *Escherichia coli* from Food-Producing Animals Slaughtered in Poland. Microb. Drug Resist. 18, 79–82. doi:10.1089/mdr.2011.0033.

Wegener, H. C. (2003). Antibiotics in animal feed and their role in resistance development. Curr. Opin. Microbiol. 6, 439–445. doi:10.1016/j.mib.2003.09.009.

Zdolec, N., Dobranić, V., Butković, I., Koturić, A., Filipović, I., and Medvid, V. (2016). Antimicrobial susceptibility of milk bacteria from healthy and drug-treated cow udder. Vet. Arh. 86, 163–172.

Zhu, Y.-G., Zhao, Y., Li, B., Huang, C.-L., Zhang, S.-Y., Yu, S., et al. (2017). Continental-scale pollution of estuaries with antibiotic resistance genes. Nat. Microbiol. 2, 16270. doi:10.1038/nmicrobiol.2016.270.

